# RAD-TGTs: Measurement of cellular tensions via flow cytometry and DNA sequencing enabled by force-dependent rupture and delivery of DNA tension probes

**DOI:** 10.1101/2022.11.08.515662

**Authors:** Matthew R. Pawlak, Adam T. Smiley, Marcus D. Kelly, Ghaidan A. Shamsan, Sarah M. Anderson, Branden A. Smeester, David A. Largaespada, David J. Odde, Wendy R. Gordon

## Abstract

Mechanical force is a key driver of cellular processes and is dysregulated in many diseases. Measuring cellular tensions to elucidate mechanotransduction pathways typically involves high-resolution but low throughput imaging of surfaces and arduous experimental preparation of materials. We present here Rupture and Deliver DNA-duplex based molecular tension sensors-RAD-TGTs. RAD-TGTs consist of immobilized DNA duplexes conjugated to a ligand and indicator (fluorophore, barcode etc) which rupture in a force-dependent manner when cells are bound. Readout of rupture is performed in cells of interest using high throughput methods such as flow cytometry and leveraging covalent DNA-protein linking HUH-tags simplifies the preparation of the tension sensor to allow use of “off-the-shelf” oligos. We demonstrate that rupture and delivery is decreased by inhibitors of cytoskeletal dynamics and knockout of mechanosensing proteins. We also show that rupture and delivery correlates with ligand affinity. Excitingly, we demonstrate that rupture and delivery of barcoded DNA-duplexes can be quantified using DNA sequencing, propelling cellular force measurements into the -omics era.

## Introduction

Mechanical force has emerged as a critical regulator of cell behavior to drive diverse biological processes from cell migration^1^ and stem cell differentiation^2^ to discrimination among similar T-cell antigens^3^. Moreover, dysregulation of cellular tensions in disease often leads to distinct mechanical phenotypes in comparison to normal cells; breast cancer cells are stiffer^4^ while metastatic breast cancer cells are more compliant that normal cells^5^. These changes in mechanical phenotype have been linked to disease progression^6^. Mechanosensing proteins in the cell sense physical stimuli in the microenvironment and convert them via protein conformational changes into a biological response in a process known as mechanotransduction. Mechanosensing pathways have become targets of mechano-therapuetics such as a growth-factor targeting antibody to reduce fibrosis in pancreatic cancer^7^. Thus identification of mechanosensing proteins that mediate disease-related mechanical phenotypes will lead to new therapeutic targets.

Piconewton forces exerted on individual proteins in the cellular context can be measured using molecular tension sensors. Numerous variations of molecular tension sensors^8,9^ have been developed in the last decade or so, including genetically-encoded FRET^9–11^ and BRET^12^ tension sensors as well as surface-immobilized peptide^13^, DNA hairpin^14,15^, and DNA duplex based tension sensors^16,17^. Molecular tension sensors are based on applied force altering the conformation of a “molecular spring” component and an optical readout of the molecular change, such as a change in fluorescence or energy transfer. Molecular tension sensors have been used to study forces sensed by mechanosensors such as integrins^18^, cadherins^19^, mucins^20^, and T-cell receptors^21^ and have begun to reveal how tensions are altered when factors in the cellular microenvironment change, such as ECM stiffness^22^ or disease-associated mutations^23^. Molecular tension sensors have massive potential to be developed into high-throughput assays that measure changes in cellular mechanical phenotype in combination with CRISPR or drug screens to drive development of the next generation of mechano-therapeutics.

The Ha lab developed DNA-based molecular tension sensors called Tension-Gauge-Tethers^16^ (TGTs), which provide an irreversible and threshold-based readout of tension. TGTs are comprised of DNA duplexes in which one strand is immobilized to a surface and other strand conjugated to a ligand recognizing a cell surface receptor. When cells are plated on top of TGTs, forces associated with adhesion and migration can mechanically rupture TGT duplexes. A fluorophore or fluorophore quencher pair incorporated into the oligos allows read out of TGT rupture via fluorescence microscopy of the surface, where gain or loss of fluorescent signal is proportional to the total number of accumulated mechanical events over time. Moreover, the tension threshold of DNA duplexes is tunable according to GC content and/or anchoring point of the bottom strand to the surface, allowing measurement of multiplexed forces. However, current versions of TGTs generally require high resolution imaging of the surface to readout duplex rupture, though a recent study reports measurement of cell-mediated forces using flow cytometry of silica beads presenting DNA tension probes ^24^. Current TGTs also require specialized glass surfaces and arduous chemical modifications of DNA to conjugate TGTs to ligands/surface, limiting their potential for high throughput assays and widespread use.

We present here “Rupture And Deliver” DNA duplex molecular tension sensors-RAD-TGTs- that allow high-throughput flow cytometry and sequencing-based measurements of duplex rupture via ligand-dependent delivery of the ruptured oligo strand into the cell of interest instead of high resolution imaging of the surface. Moreover, RAD-TGTs can be prepared simply with “off the shelf” oligos by leveraging covalent DNA-linking HUH-tags^25^ to attach desired recombinant protein ligands of interest to unmodified DNA. RAD-TGTs can be plated in commercial 96 well plates with minimal surface preparation. We demonstrate that RAD-TGTs provide a relative measure of cellular forces, with a signal proportional ligand affinity. We validate that internalized fluorescence is sensitive to cytoskeletal modulators as well as CRISPR knockout of relevant cellular mechanosensors. Finally, we show for the first time that cellular forces can be measured via the sequence of internalized DNA-based molecular tension sensors, greatly expanding the potential for multiplexed, high-throughput assays.

## Results

### RAD-TGTs: Design of “off-the-shelf” Rupture and Deliver DNA tension probes leveraging HUH-endonucleases

The overall goal of this study was to develop a high-throughput assay to measure the cumulative tension a cell exerts onto its environment. We reasoned that existing DNA-duplex molecular tension probes, also known as Tension Gauge Tethers (TGTs), could be converted into a high-throughput platform by changing the focus of the readout from the oligo remaining on the surface to the oligo captured by the cell upon rupture (**Fig 1a**). We hypothesized that tension dependent rupture of TGTs and subsequent delivery of the ruptured oligo into the cell of interest could be detected by flow cytometry or Next Generation Sequencing. RAD-TGTs were adapted from the TGT designs originally reported by the Ha group^26^ which consist of two strands of DNA one that is conjugated to a ligand (the ligand strand) and a strand which contains a modification to immobilize the duplex (the anchor strand) (**Fig 1b)**. The duplex component of the TGT remains identical to previous designs so as not to alter the thermodynamic properties of DNA duplex formation. The anchor strand contains a biotin modification either at the 5’ or 3’ end that binds plated neutravidin to yield TGTs that rupture in the unzipping (12pN) and shear (54pN) modes, respectively.

**Figure 1.**
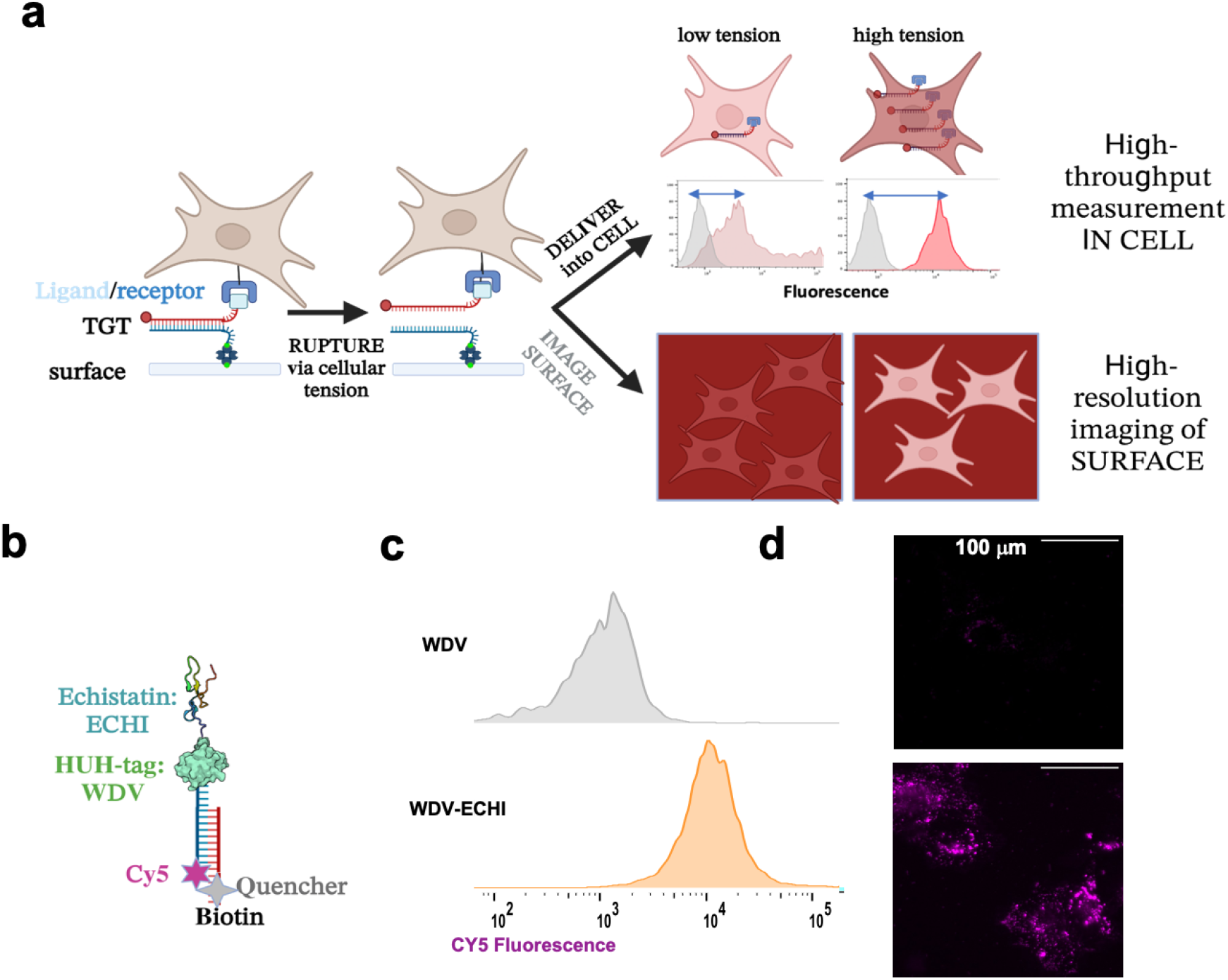
Rupture and Deliver Tension-Gauge-Tethers: RAD-TGTs. **a.** Schematic of concept of RAD-TGTs to readout rupture of TGTs in the cell of interest instead of high-resolution surface imaging. **b**. Overall design features of RAD-TGTs **c.** Histograms of CY5 fluorescence from of CHO-K1 cells adhered to RAD-TGTs tethered to WDV or WDV-echistatin integrin ligand **d.** qTGT surface fluorescence imaging readout analogous to **c,** with Cy5/Quencher swapped compared to **c** such that TGT rupture turns on fluorescence.

Typically, the ligand strand of TGTs is chemically modified with a ligand such as integrin-binding cyclic-RGD using amino, thiol, or Click chemistry. Our design leverages DNA-to-protein linking HUH endonucleases to allow covalent attachment of versatile protein based ligands to RAD-TGTs. Briefly, HUH-endonucleases are small (<40 kDa) proteins that can be fused to ligands of interest to form robust, sequence-specific phospho-tyrosine covalent bonds with a short nona-nucleotide sequence of ssDNA within minutes under physiologic conditions^25^. Thus the ligand strand does not require chemical modification as in other TGT designs, but simply includes a short 5’ DNA extension. For readout of rupture and delivery into cells, the ligand strand also contains a 3’ fluorophore or a short nucleotide barcode. We also include a quencher on the anchor strand to ensure detection of ruptured oligos and not spurious detachment of duplex DNA during cell handling.

The final component of RAD-TGT design is the choice of ligand for a high-throughput readout. The majority of studies use integrin-binding linear or cyclic RGD peptides due to the relative ease of chemical conjugation to DNA. These peptides have a broad range of affinities (low nM to μM)^27^ for different integrin subtypes and result in an estimated 1000-10,000 rupture events per cell^28^, which typically requires high NA objectives and high sensitivity cameras to image. To ensure robust signal per cell for downstream flow cytometry readout, we chose to use integrin-binding echistatin, a small toxin-derived protein presenting the RGD peptide in a tight loop with subnanomolar affinity for every subtype of integrin tested ^27^. Echistatin is fused to an HUH-tag called WDV^29,30^ and is covalently tethered to annealed duplex TGTs prior to immobilization (Supplementary Fig.1). A WDV only TGT control is always used for background correction.

### Flow cytometry detects rupture and delivery of immobilized RAD-TGTs

We first assembled immobilized RAD-TGTs (**Fig 1b**) to determine if internalized fluorescence derived from TGT rupture could be detected by flow cytometry. We attached WDV and WDV-echistatin ligands to duplex TGTs and immobilized them. Briefly, assembly of the RAD-TGTs (see Methods for full details) involves: 1) coating standard glass bottom 96 well plates with biotin-BSA, 2) incubation with neutravidin, and 3) incubation with 80ul of a 1uM solution of assembled TGTs overnight. After washing, approximately 15,000 cells per well are plated in serum free media to mitigate DNA degradation by nucleases for 1-4 hours. At the desired timepoint, the cells are trypsinized and analyzed by flow cytometry.

We first tested RAD-TGTs on CHO-K1 cells, commonly used in TGT experiments. Flow cytometry of the CHO-K1 cells 2 hours after plating on RAD-TGTs shows a clear difference between the negative control WDV and WDV-echistatin, with the fold change of median fluorescence being about 10-fold **(Fig 1c)**. Two distinct and relatively narrow populations are observed, in contrast to other studies detecting cellular forces by flow cytometry of microparticles containing DNA tension probes, which instead report a modest broadening of the population in response to cellular tension^24^. This suggests that a homogeneous population of cells plated on the RAD-TGTs are exerting tension on the surface. We performed a parallel readout of qTGT rupture by surface fluorescence microscopy **(Fig 1d)** to validate our surface and TGT preps and confirm the flow cytometry results, with the Cy5 fluorophore on the bottom strand and the quencher on the top strand to allow a gain of fluorescence readout when the RAD-TGT ruptures. The CHO-K1 cells plated on WDV-Echi show a strong fluorescence footprint under the cells while WDV alone shows very little fluorescence. Additional characterization and optimization of the RAD-TGT protocol confirmed that internalized fluorescence increases with time after cell plating up to about 2 hours (Supplementary Fig.2) and that overall median fluorescence does not depend strongly on cell density (Supplementary Fig.3). We also performed measurements including fibronectin in the coating protocol or on surfaces with mixed 12pN and 54 pN TGTs to ensure the results are not confounded by differences in cell behavior on 12pN TGTs^31^ (Supplementary Figs. 4 and 5). Generally, all surface preps result in the same trends in the data and cells adhering well after 2 hours with varying morphologies as expected (Supplementary Fig.6). Initial attempts using commercial polystyrene neutravidin coated plates resulted in similar trends in median fluorescence as glass bottom plates, but an overall broader distribution of fluorescence (Supplementary Fig.7) which we attributed to non-specific adsorption to polystyrene. Thus we proceeded using glass bottom plates, though it should be noted that our prep does not involve any special cleaning or silanization of the surface as many other protocols do.

### RAD-TGT signal is modulated by cytoskeletal inhibitors and loss of putative mechanosensors

To gain more insight into the role of cellular forces on the observed fluorescence signal, we next measured effects of cytoskeletal modulators on delivery of the ruptured oligo into the cell. We also expanded the studies to U251 glioma cells. We observed that U251 cells exhibit an enhanced fold change in median fluorescence compared to CHO-K1 cells **(Fig 2a,b)**, suggesting they exert higher cellular forces. Indeed, an echistatin titration of integrins on U251 and CHO-K1 cells revealed an 3-4 fold difference in bound echistatin, suggesting a greater number of integrin clutches available to exert traction force (Supplementary Fig.8). We also plated CHO-K1 and U251 cells on mixed RAD-TGTs with intrinsically different tension tolerances (**Fig 2c**). A mixture of Cy5 labeled 12pN duplex and Alexa488 labeled 54pN duplex resulted in distinct populations on a 2D dot plot, demonstrating that RAD-TGTs can be multiplexed to add additional parameters to the mechanical signature that could be useful in high throughput screening and flow sorting of cells exhibiting distinct mechanical signatures. Generally, the 54pN RAD-TGTs resulted in similar trends as the 12pN tension tolerance. We next treated cells with Myosin II inhibitor para-amino-Blebbistatin **(Fig 2a,b)**, which has been shown to decrease TGT rupture in CHO-K1 cells ^32^ and decrease traction force in U251 cells ^33^. Indeed para-amino-Blebbistatin treatment resulted in a consistent ~30% decrease in fluorescence in both cell lines **(Fig 2b)**. These results suggest that the observed fluorescence signal is related to the traction force the cell is exerting.

**Figure 2.**
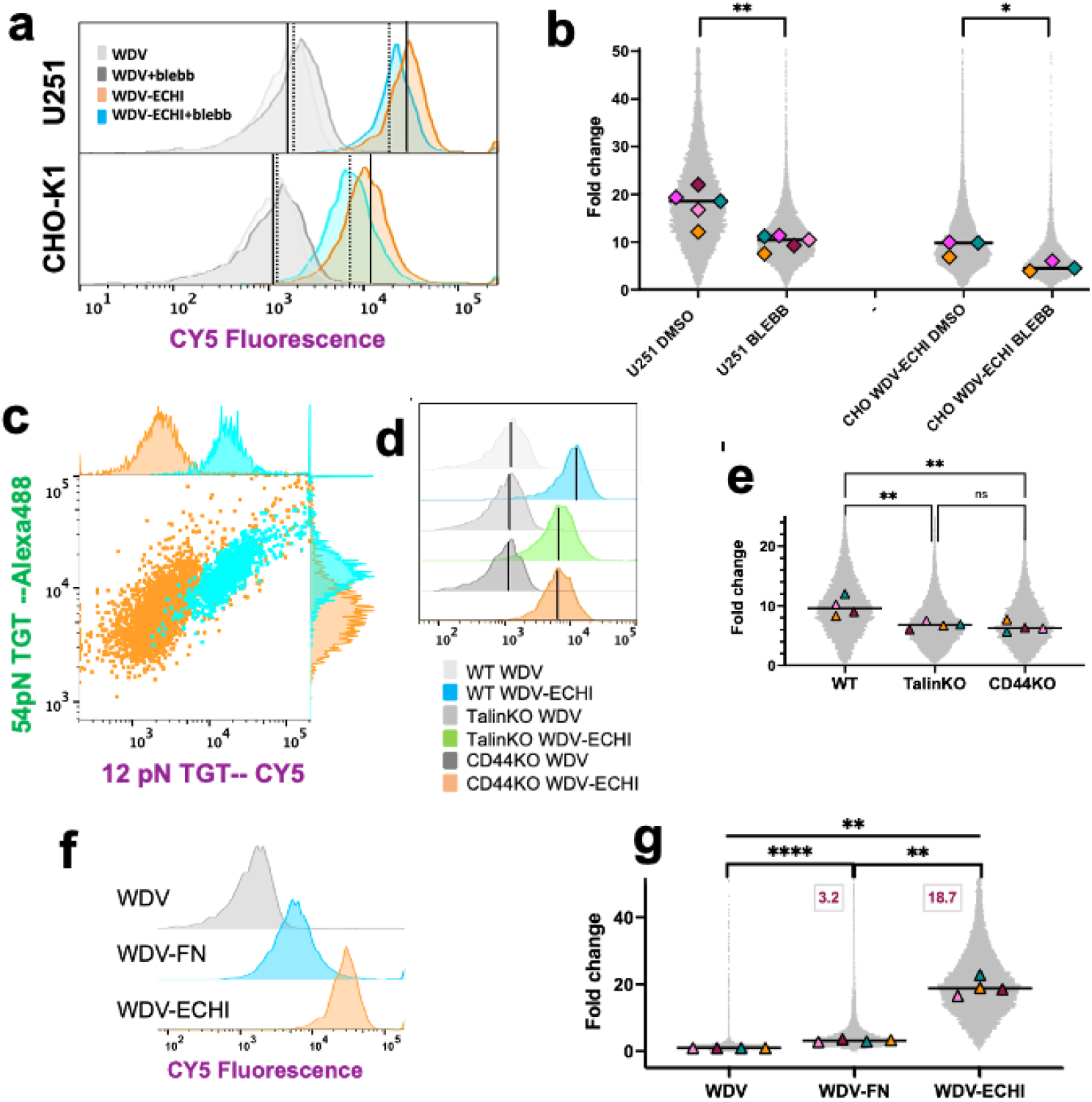
Rupture and Deliver TGTs provide a relative measure of cellular traction force and ligand affinity. **a,b** Myosin II inhibitor para-amino blebbistatin (blebb) reduces rupture and delivery in CHO-K1 and U251 cells. **a.** Representative histograms of DMSO and Blebb treatment of cells adhered to WDV and WDV-ECHI RAD-TGTs. Black solid and dashed lines denote median of DMSO and blebb treatments, respectively **b**. Superplots of biological replicates of median fluorescence fold changes upon drug treatments from a. Medians of the biological replicates (diamonds) are overlaid on dot plots of all of the data normalized to the corresponding WDV alone median. **- 0.0061; *- 0.0364 **c.** RAD-TGTs can be multiplexxed to form 2D plots. Cells plated on surfaces containing CY5 labelled 12 pN RAD-TGTs and Alexa488 labelled 54 pN RAD-TGTs all TGTs contained WDV-Echi. CHO-K1 cells (orange) and U251 cells (cyan) are shown as 2D Alexa488 versus Cy5 dot plots. It should be noted that the bottom strand quencher was not present in this experiment, inclusion would decrease noise allowing for greater separation of populations. **d,e.** Talin1 and CD44 CRISPR knockouts in U251 cells reduce rupture and delivery of RAD-TGTs. **d**. Representative histograms of U251, U251-talinKO and U251-CD44KO cells plated on WDV and WDV-echistatin RAD-TGTs, black line denotes median fluorescence. **e**. Superplots of biological replicates of data shown in c. as described in b. **talinKO-0.0087; **CD44KO-0.005; ns-0.9243 **f,g.** Rupture and delivery depends on ligand affinity. **f**. Representative histogram of U251 cells plated on WDV, WDV-FN, or WDV-echistatin RAD-TGTs. **g**. Superplots as described above. ****<0.0001; **FN/echi-0.0057; **wdv/echi-0.0083 : All statistics were performed using ANOVA of the medians of biological replicates as previously described^42^.

The high throughput readout of RAD-TGT rupture could enable rapid screening for knockout of cellular proteins that alter cellular force generation. As proof of concept, we compared wild type U251 cells (WT) against U251 cells with CRISPR knockouts of two mechanosensing proteins-talin-1 and CD44 **(Fig 2d,e)**. Talin-1 is a mechanosensor in the integrin-anchored focal adhesion complex while CD44 displays mechanosensitive behavior in glioma cells^34^. U251 talinKO cells displayed a rounded morphology on TGTs, while CD44 cells resembled wildtype U251 cells (Supplementary Fig 6). Interestingly, RAD-TGT analysis of all three U251 cell lines (WT, TLN1 KO, CD44 KO) revealed statistically significant decreases in signal from both KO’s relative to WT. These results suggest that KO’s of these genes decreased cellular force generation compared to wildtype cells, underscoring that we are observing changes in traction force while also providing proof-of-concept for the use of RAD-TGTs in high throughput CRISPR screens.

### RAD-TGT can discriminate among ligands of different affinity

We hypothesized that the rupture and delivery readout would be sensitive to the affinity of the ligand-receptor interaction, as the rupture of the DNA duplex depends not only on the tension generated in a ligand-receptor complex but also on the kinetics of the ligand-receptor interaction. Echistatin is known to have a sub-nM affinity for a broad range of integrin receptors^27^, increasing the propensity to rupture TGTs. We thus made HUH fusions with a chimeric fibronectin domain (FN) predicted to have more typical affinity to integrins and facilitate optimal cell migration ^35^ We indeed observed dramatic differences in median fluorescence between the two ligands **(Fig 2f,g)** that were detectable over background fluorescence (Supplementary Fig.9). Though experiments with the FN ligand produced a lower fold change from the WDV negative control compared to echistain, all of the trends observed with echistatin were recapitulated with the FN ligand (Supplementary Fig.10).

### RAD-TGT rupture can be measured by DNA-sequencing

We envisioned that the multiplexability of RAD-TGTs could be expanded beyond the color barrier of flow cytometry using DNA sequencing methods such as Next Generation Sequencing. As proof of concept, we barcoded RAD-TGTs conjugated to three different ligands HUH, HUH-FN, and HUH-echistatin, and immobilized them together. The barcode sequence was added to the 3’ end of the ligand strand. We performed Sanger sequencing (**Fig 3a**) and Illumina Next Generation Sequencing (NGS) (**Fig 3b)** of the amplified cell lysate. We used web-based sequence deconvolution programs to calculate the percent of each barcode represented in the samples quantified by sanger sequencing and Biopython to quantify NGS data. Interestingly, we observed a statistically significant increase in HUH-echistatin relative to HUH and HUH-FN with both sequencing methods but we only saw a statistically significant increase in HUH-FN with sanger sequencing. It should be noted that when the experiment was performed with fluorescent oligos and a flow cytometry readout the results mirrored the NGS data in which only echistatin was elevated (Supplementary Fig. 11). We attribute the robust echistatin signal relative to other ligands to the very strong affinity of echistatin for integrins. We attribute the discrepancies between Sanger sequencing and NGS to implicit error rates of sequencing that can arise from events such as PCR bias, future work to improve this may include the addition of multiple barcodes per condition to mitigate any bias.

**Figure 3.**
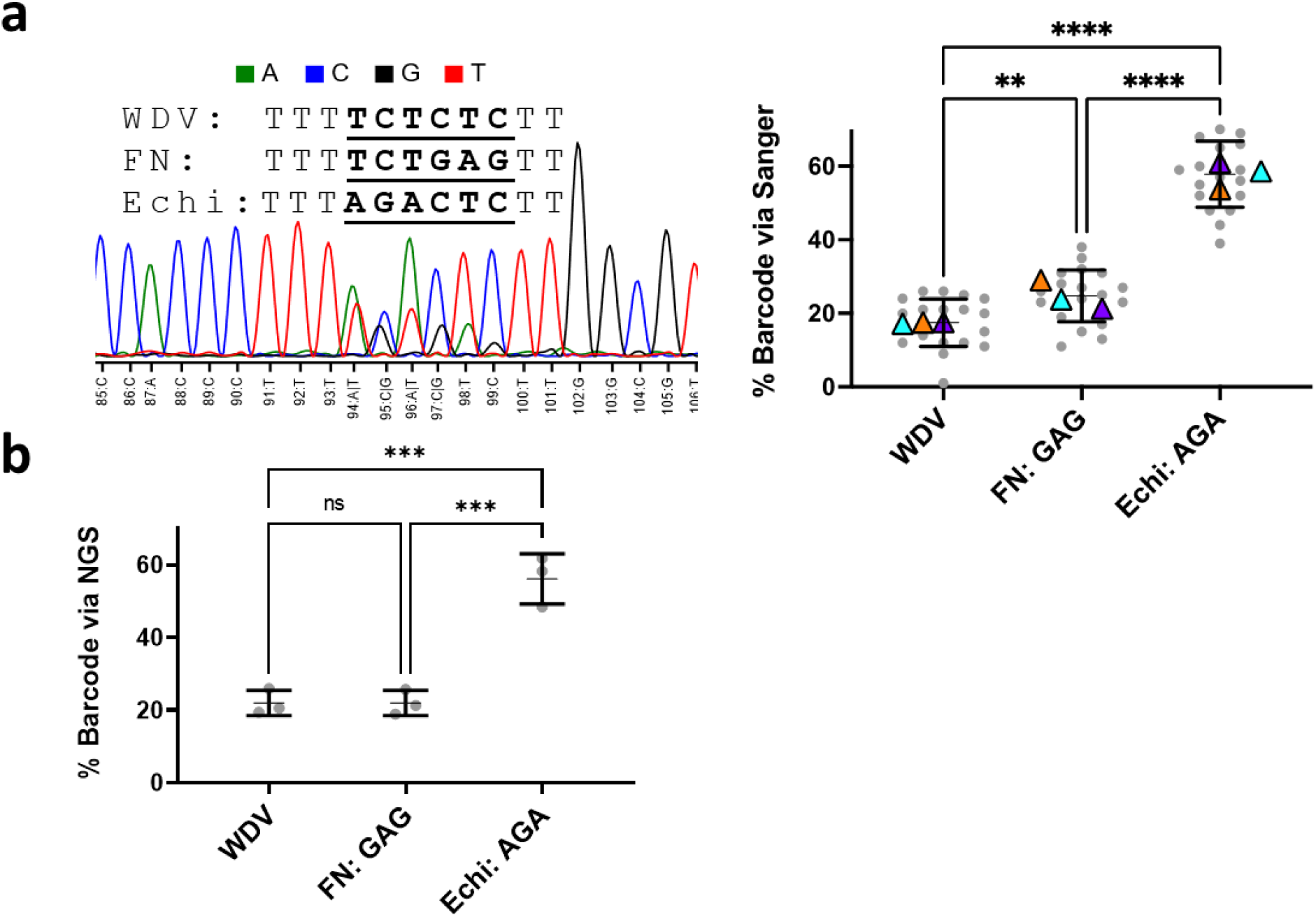
Cellular tension can be measured by DNA sequencing of barcoded RAD-TGTs. RAD-TGTs were conjugated to WDV, WDV-FN, and WDV-ECHI, mixed 1:1:1 at ⅓ standard concentration and immobilized. U251 cells were plated, trypsinized and lysed. PCR of the barcodes of three replicates was performed and sent for forward and reverse Illumina and Sanger sequencing. Deconvolution analysis was performed as described in the methods. **a.** Representative Sanger sequence (left) and analysis of the data (right). Mean % barcode for each replicate is denoted by the triangles, the center line within standard deviation bars is the total mean % barcode. **wdv/fn-0.0077; **** fn/echi<0.0001; ****wdv/echi<0.0001. **b.** Analysis of NGS sequencing measuring the percentage of total reads each barcode composed, following the same convention as a. Each replicate contains a singular data point so no triangles are present. ***p<0.001.

## DISCUSSION

The ability to quantify cellular forces using methods such as traction force microscopy (TFM) and molecular tension sensors has established mechanical force as a critical regulator of cellular processes and demonstrated drastic differences in mechanical phenotype between normal and diseased cells^6^. However, the low throughput nature of reading out forces by high-resolution microscopy as well as complex preparation of elastic surfaces and functionalized DNA-based tension sensors have precluded studies of the mechanosome from entering the -omics era. The Rupture and Deliver DNA-based molecular tension sensors (RAD-TGTs) presented here offer a useful tool to close this gap.

The major advance offered by RAD-TGTs is the readout of immobilized DNA duplex rupture by cells of interest via delivery of ruptured oligos into cells using high throughput readouts. We demonstrated that flow cytometry provides a sensitive method to readout ruptured and internalized fluorescent oligos of thousands of cells in minutes in one to three channels, but could be expanded to additional colors. The readout was sensitive enough to detect oligos conjugated to a single fluorophore in multiple cell lines without requiring additional amplification steps used in recent studies^28^. However the sensitivity varies among cell lines, depending on adhesive and traction forces generated, thus the flow cytometry readout may not be suitable for cell lines that generate low traction force. The sensitivity of the flow based readout could be improved by attaching tandem fluorophores or even quantum dots. Recently, the Salaita group developed silica-bead-presenting TGTs that fluoresce following rupture^24^. Beads are added to cells and analyzed using flow cytometry which allows for high throughput measurement. However, the overall fold change of fluorescence was significantly lower than RAD-TGTs and analyzing beads alone cannot resolve what specific cell was encountered thus preventing use in heterogeneous populations. RAD-TGTs resolve this difficulty by directly measuring individual cells. This further highlights the utility of this sensor in applications such as CRISPR-KO screens and studying heterogeneous environments such as the tumor microenvironment.

Excitingly, we demonstrated that the ruptured DNA oligo could also be detected by sequencing captured barcodes in cell lysates. We demonstrated proof of concept detection of three barcoded RAD-TGTs harboring different ligands, via Sanger sequencing and NGS. The ability to sequence captured barcodes in combination with NGS could permit further multiplexing of ligands and TGTs of variable tension tolerances in the context of large drug or CRISPR screens. This could be used to screen large libraries of ligands recognizing patient T-cells or for whole genome CRISPR-KO screens to identify key mechanosensors underlying a given mechanical phenotype.

The use of HUH-endonucleases in RAD-TGTs expands the repertoire of proteins that can be attached to TGTs to allow a probing a broader range of receptor-ligand interactions, adaptation to more complex microenvironments such as 3D hydrogels, and expansion of readouts possible after rupture and delivery. For example, the “anchor” strand of TGTs could be conjugated via HUH-tags to collagen binding CNA35^36^ to allow immobilization of TGTs in collagen gels. The ligand strand could be conjugated to luciferases, nanobodies, or even genome engineering reagents to allow readout of TGT rupture in animals or tension-dependent delivery of reagents. Here, the use of the integrin binding protein-based ligands, such as broad spectrum echistatin with sub-nM affinity enabled flow cytometry readout via increased rupture events detected per cell. Moreover, because the HUH-ligand is added post-duplex assembly, longer-term studies of cellular tension may be possible by adding HUH-ligands at variable timepoints after plating on unreactive RAD-TGTs.

RAD-TGTs provide a relative measure of cellular forces, akin to traction force microscopy. We demonstrated that treatment with cytoskeletal modulating drugs such as para-amino-Blebbistatin reproducibly reduced the levels of fluorescence. Moreover, knockout of the key focal adhesion mechanosensor talin and putative hyaluronan mechanosensor CD44 also reduced fluorescence. TFM provides an absolute measure of traction forces, but relies on large enough forces to perturb beads in an elastic gel and typically involves measurement of tens of cells over hours compared to thousands of cells in minutes. Moreover, elastic substrates can be challenging to prepare in 96-well plates and analysis can be difficult and variable among studies. While molecular tension sensors cannot provide absolute quantification of traction forces because it relies on stochastic ligand-receptor interactions and thus does not capture all events, these sensors will be useful for comparisons of relative cellular tensions among multiple conditions or treatments.

Finally, the RAD-TGTs presented in this study were all assembled using oligos purchased from IDT, with no additional chemical modification, and ligands containing HUH-fusions can be easily prepared using *E. coli* expression protocols. Moreover, commercial 96 well plates were used with minimal surface preparation. Thus the advances offered by RAD-TGTs should make DNA-tension sensor probes accessible to more research labs.

## Methods

### HUH-Ligand Preparation

HUH variants were expressed in *E. coli* and purified via subsequent Ni-NTA affinity chromatography and size exclusion chromatography following previously described protocols^25,29^.

### RAD-TGT Synthesis

TGTs were designed using previously characterized DNA sequences^26^ with a 5’ extension to allow for HUH binding. The following oligonucleotides were purchased from Integrated DNA Technologies

Fluorescently Labeled Ligand Strand: 5-GCT ATA AAC TCA CCG TAA TTT TTT GGC CCG CAG CGA CCA CCC TTT /3Cy5Sp/-3

Quencher Ligand Strand: 5-GCT ATA AAC TCA CCG TAA TTT TTT GGC CCG CAG CGA CCA CCC TTT/3IAbRQSp/-3

Non-Anchor Strand: 5-GGG TGG TCG CTG CGG GCC-3

12 pN Quencher Anchor Strand: 5-/5IAbRQ/GGG TGG TCG CTG CGG GCC/3Bio/-3

12 pN Unlabelled Anchor Strand: 5-GGG TGG TCG CTG CGG GCC /3Bio/-3

12 pN Fluorescently Labeled Anchor Strand: 5-/5Cy5/GGG TGG TCG CTG CGG GCC /3Bio/-3

54 pN Quencher Anchor Strand: 5- /5IAbRQ/iBiodT/GGG TGG TCG CTG CGG GCC-3

Barcoded Ligand Strand 1: 5-GCT ATA AAC TCA CCG TAA TTT TTT GGC CCG CAG CGA CCA CCC TTT TCT CTC TT GGC GTC ATC GTG TAC CGG-3

Barcoded Ligand Strand 2: 5-GCT ATA AAC TCA CCG TAA TTT TTT GGC CCG CAG CGA CCA CCC TTT AGA CTC TT GGC GTC ATC GTG TAC CGG-3

Barcoded Ligand Strand 3: 5- GCT ATA AAC TCA CCG TAA TTT TTT GGC CCG CAG CGA CCA CCC TTT TCT GAG TT GGC GTC ATC GTG TAC CGG-3

Forward Amplicon Primer: 5-ACA CTC TTT CCC TAC ACG ACG CTC TTC CGA TCT CTC ACC ATG TGG TGA CGC CAG AAT TTG CCG CAA TAC ACA GTT TAC GCC GTT CGG TCA GCT TGG TAT CCG TAG CGC AGC GAC CAC CCT TT-3

Reverse Amplicon Primer: 5--GAC TGG AGT TCA GAC GTG TGC TCT TCC GAT CTG AGA AAC CTA AGC AGA CTT CTC CTG GTC GAT GAT TGA TAA GGG TCT CGG AAT GTC CCC GGT CGC ATG GTT CCG GTA CAC GAT GAC GCC-3

To prepare TGTs, anchor and ligand strands were mixed in a 1.1:1 ratio and then annealed in a 1x annealing buffer (10 mM Tris pH 7.5, 50 mM NaCl, 1 mM EDTA) by heating at 98 c for 5 minutes followed by cooling at room temperature for 1 hour. Excess bottom strand was used to mitigate any single stranded fluorescent top strand which may be internalized resulting in false positives.

RAD-TGTs were then generated by reacting HUH-ligand of interest with the annealed duplex in a 2:1 ratio. Reactions were performed in the following buffer (50 mM HEPES pH 8.0, 50 mM NaCl, 1 mM MnCl_2_) over 30 minutes at 37° C.

### Surface Preparation

96 well glass plates (Matek, PBK96G-1.5-5-F) were incubated with 80 μL of 100 ug/ml BSA-Biotin (Thermo Fisher, 29130) in PBS for 2 hours at room temperature. In specified contexts, 18.75 ug/ml of fibronectin was added to the BSA-biotin solution. Wells were rinsed 2x with cold PBS and incubated with 100 ug/ml neutravidin (Thermo Fisher, 31000) for 30 minutes at room temperature. Wells were rinsed once more and incubated with 80 μL of 1 μM RAD-TGT and were incubated at 4° C overnight. It is essential that the wells never dry to prevent any unintentional adsorption to the surface, for all experiments volume in the well never fell below 50 μL.

### Cell lines

U251 glioma cells were a gift from the Odde lab and CHO-K1 cells were purchased from ATCC (ATCC, CCL-61). U251 talin-KO and U251 CD-44 KO cells were prepared using published methods ^37,38^. TLN1 KO and CD44 KO were achieved using the CRISPR/Cas9 system. A guide RNA (sequence AACUGUGAAGACGAUCAUGG) was created to target TLN1. A guide RNA (gRNA) was created to target exon 2 in CD44 human cell lines (GAATACACCTGCAAAGCGGC). A co-transposition method was used to enhance screening for knockout clones. Briefly, cells were transfected with Cas9 nuclease, gRNA, PiggyBac transposases, and PiggyBac transposon plasmid containing puromycin selection using FuGENE (Promega, Madison, WI) following the manufacturer’s protocol. Puromycin selection was performed and single cell clones were generated using serial dilution. After transfections the cells were split into single clones and western blot was used to confirm the knockout. WB also verified that there was not over expression of Talin2 in response to KO of Talin1.

### Cell Culture

Cells were maintained for no more than 15 passages. All cells were regularly passaged every 2 to 3 days when 80% confluency was achieved. All cells, other than CHO-K1 cells, were maintained in Dulbecco’s Modified Eagle Medium (Corning, 10027CV) supplemented with 10% fetal bovine serum (FBS) (R&D Systems, S11150) and penicillin/streptomycin (Gibco,15070063). CHO-K1 cells were maintained in F-12K medium (ATCC, 302004) supplemented with 10% FBS and penicillin/streptomycin.

### RAD-TGT Experimental Setup

Cells of interest were trypsinized (Gibco, 25200056) for 5 minutes and transferred to Opti-MEM Reduced Serum Media (Gibco, 31985062) following this cells were counted using an automated cell counter (Countess II, Invitrogen). If cells were to be treated with para-Amino-Blebbistatin (Cayman Chemical, 22699), the cells were incubated with 50 μM para-Amino-Blebbistatin at 37c for 30 minutes before application to the RAD-TGT surface. All para-amino-Blebbistatin experiments were accompanied by a vehicle control consisting of cells being incubated in an equivalent volume of DMSO (Invitrogen, D12345). RAD-TGT surfaces were washed once with cold PBS and 2x with Opti-MEM. Following washes and any drug treatment 15,000 cells were added to the wells and incubated at 37c with 5% CO_2_ for desired period of time, typically 2 hours.

### Flow Cytometry Experiments

Following incubation on the RAD-TGT medium was removed and 30 μL of trypsin was added to each well. The plate was further incubated at 37° C for 5 minutes. Following trypsinization 170 μL of flow cytometry buffer (PBS with 1% FBS and 1 mM EDTA) was added to the wells to quench the trypsin and resuspend the cells. Cells were removed from wells and analyzed with a BD Accuri C6 Plus Personal Flow Cytometer without further washing. Cells were gated from the collected data, followed by gating for individual cells and then gated for cells that had a detectable signal.

### Sequencing Experiments

Sequencing experiments followed the same method as flow cytometry experiments but cells were not analyzed with a cytometer. Three unique RAD-TGTs were in the experimental well, one for each ligand (WDV, FN, or Echi) and each RAD-TGT contained a novel barcode on the 3’ end of the ligand strand. Following removal from the well, cells were treated with 1x Passive Lysis Buffer (Promega, E1941) for 20 minutes at room temperature while on an orbital shaker. The lysate had the barcoded top strands PCR amplified and the resulting PCR product was gel purified and submitted for Sanger and Illumina Sequencing. Quantification of barcodes in the Sanger sequences was performed using the Base-editing analysis software EditR^39^. A dummy guide sequence was denoted that spanned the barcode area. Percentages at AGA and GAG in the barcodes were averaged to calculate percent of Echi and FN respectively for forward and reverse sequencings of three biological replicates. WDV was calculated from the difference from 100%. The program Tracy^40^ was used to visualize the sequence displayed in Fig 3. Quantification of barcodes in the Next-Generation sequencing assay was performed with a custom Python script using the Biopython package^41^. Briefly, forward sequencing reads were parsed, trimmed to include only the region containing the barcode, and barcodes were then counted. Reverse sequencing reads were reverse complemented and then processed in an identical manner. Forward and reverse counts for the three barcodes in each sample were combined and barcodes corresponding to either Echi, FN, or WDV were exported for each sample.

### Microscopy

To visualize TGT ruptures surfaces were prepared following the above protocols. TGTs consisted of Quencher Ligand Strand and a Fluorescently Labeled Anchor Strand so if ruptured one should observe a gain of fluorescence on the glass surface. Rather than dissociating the cells post trypsinization, each well had the media removed and Fluorobrite DMEM (Gibco, A1896701) was immediately added to wells. Cells were then imaged at 40x with an EVOS FL AUTO fluorescent microscope. Images were processed with ImageJ

### Statistical Analysis

In general, each experiment was performed on at least three separate days with 1-3 separated wells per day. The superplots were generated by importing histogram data from FlowJo into Prism. The ligand data were normalized to the median of the WDV alone data. The fluorescence data were concatenated to produce the dot plots. The medians of the normalized data for each biological replicate were used to overlay on the dot plot, and statistical significance was calculated using unpaired ANOVA as previously described^42^.

## Acknowledgements

We acknowledge funding from the NIH R35GM119483 and U54210190. WRG is a Pew Biomedical Scholar. M.D.K. acknowledges a Cancer Center Training grant (T32CA009138).

## Author contributions

MRP performed the majority of the experiments in this study, analyzed data, and wrote the manuscript. ATS performed Sanger and NGS sequencing experiments. MDK performed experiments related to this study. WRG analyzed data and wrote the manuscript. GAS, SMA, BAS, DAL, DJO engineered the CRISPR KO cell lines used in this study.

## Supplementary Figures

**Supplementary Fig. 1.**
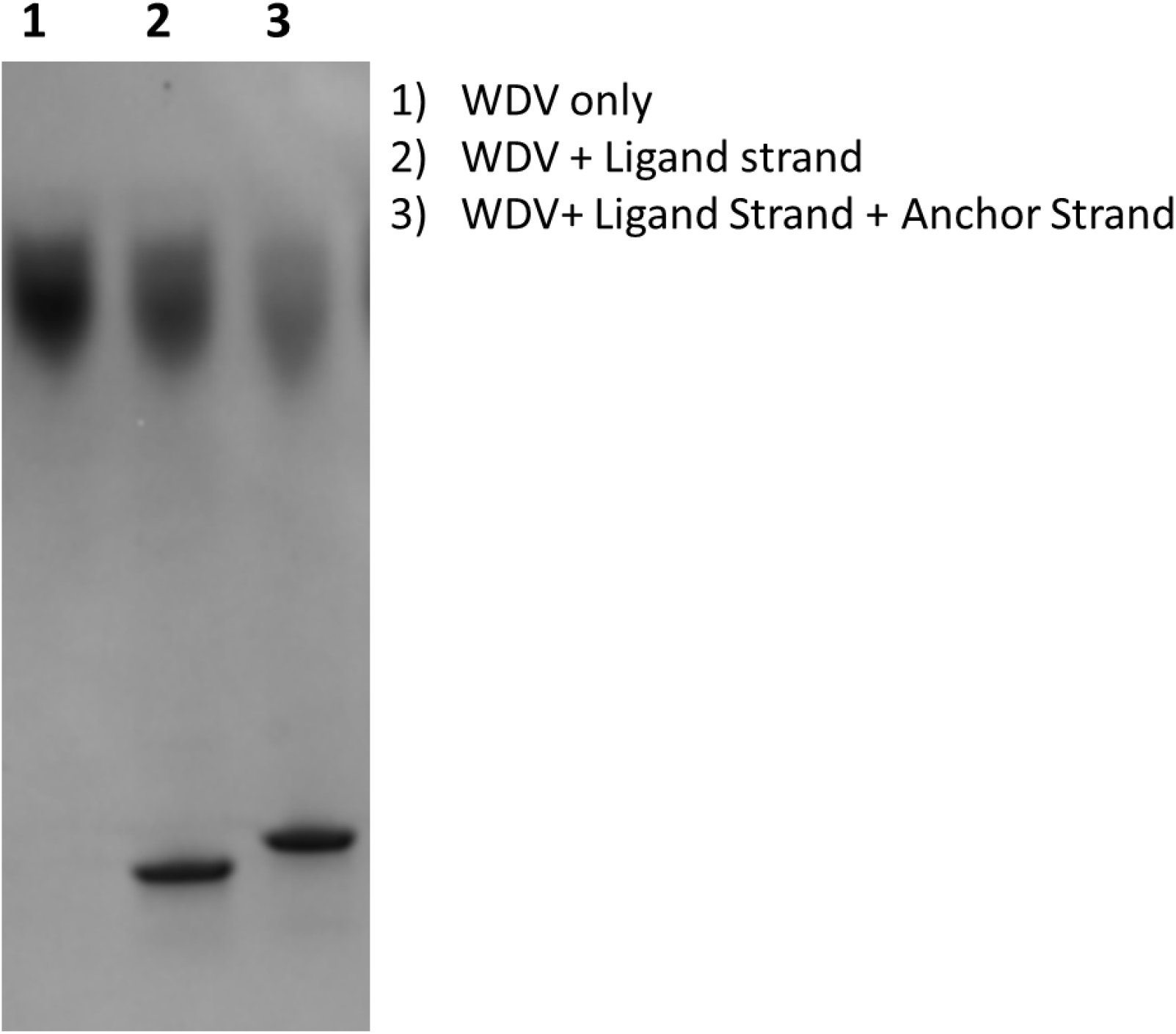
HUH reacting with ssDNA and duplex DNA captured via native gel electrophoresis. RAD-TGT components were reacted and analyzed in a non-denaturing polyacrylamide gel. The reaction for lane 1 only contained WDV, lane 2 contained the ligand strand from RAD-TGT and excess WDV, lane 3 contained both ligand and anchor strands from the RAD-TGT annealed together and excess WDV. Lane 1 contains a singular band representing WDV while lanes 2 and 3 contain two bands, WDV and a WDV-DNA band below that. The new bands in lanes 2 and 3 is the WDV reacting with the oligos present, the oligos carry a negative charge so when bound to WDV migration distance increases on a native gel. The band in lane 3 is slightly shifted up relative to the band in lane 2, this is because the annealed duplex increases the mass of the complex resulting in slower migration. From this gel it is evident that the WDV can react with the duplex DNA present in RAD-TGTs.

**Supplementary Fig. 2.**
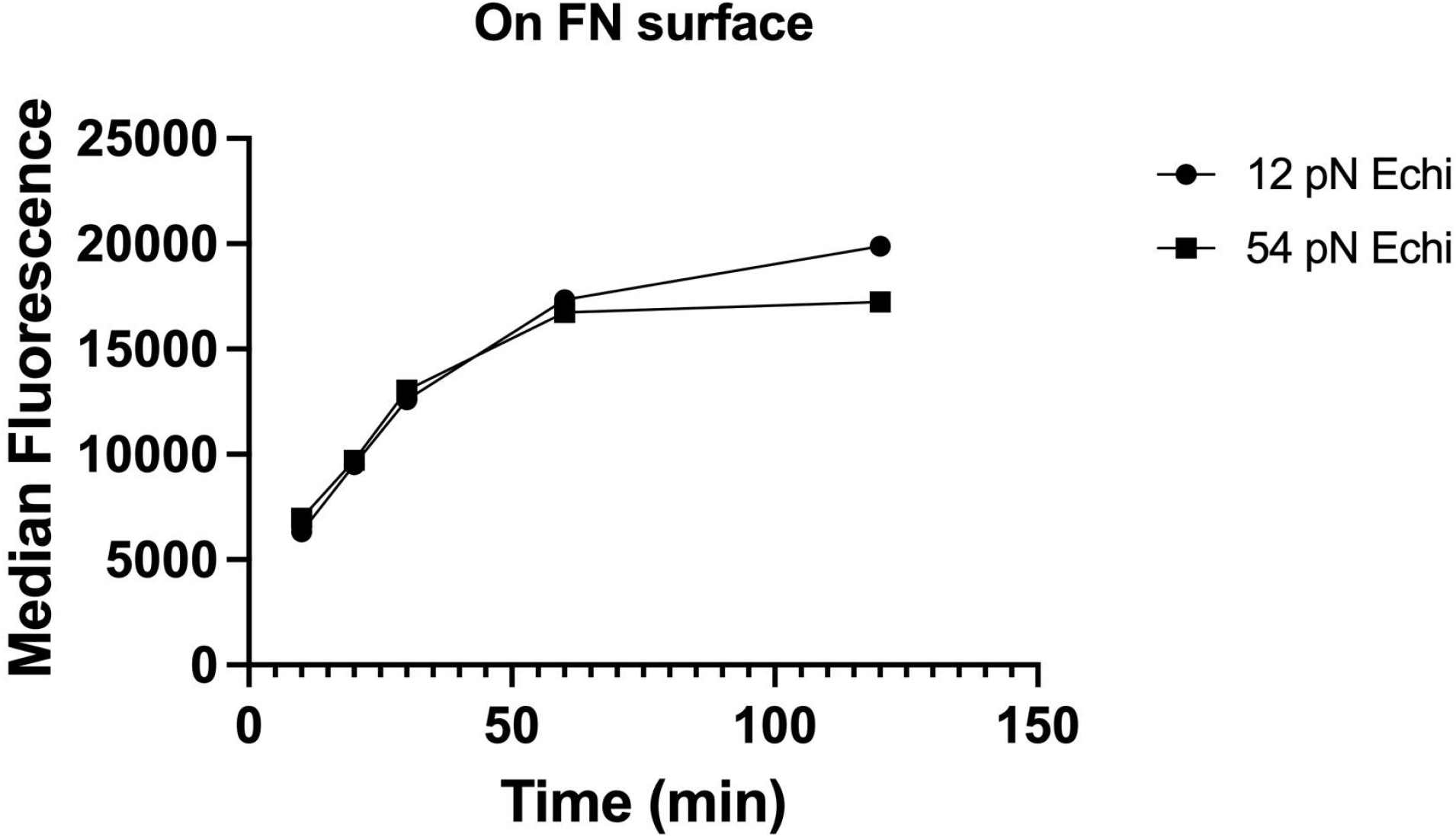
Incubation time on RAD-TGTs increases fluorescent intensity. U251 cells were incubated on an echistatin conjugated RAD-TGT surface that also was coated with fibronectin. Cells were incubated for varying time points from 10 minutes to 120 minutes, once desired time was reached cells were dissociated and immediately analyzed via flow cytometry. This was performed with both 12 and 54 pN RAD-TGTs and a clear increase in signal over time was observed.

**Supplementary Fig. 3.**
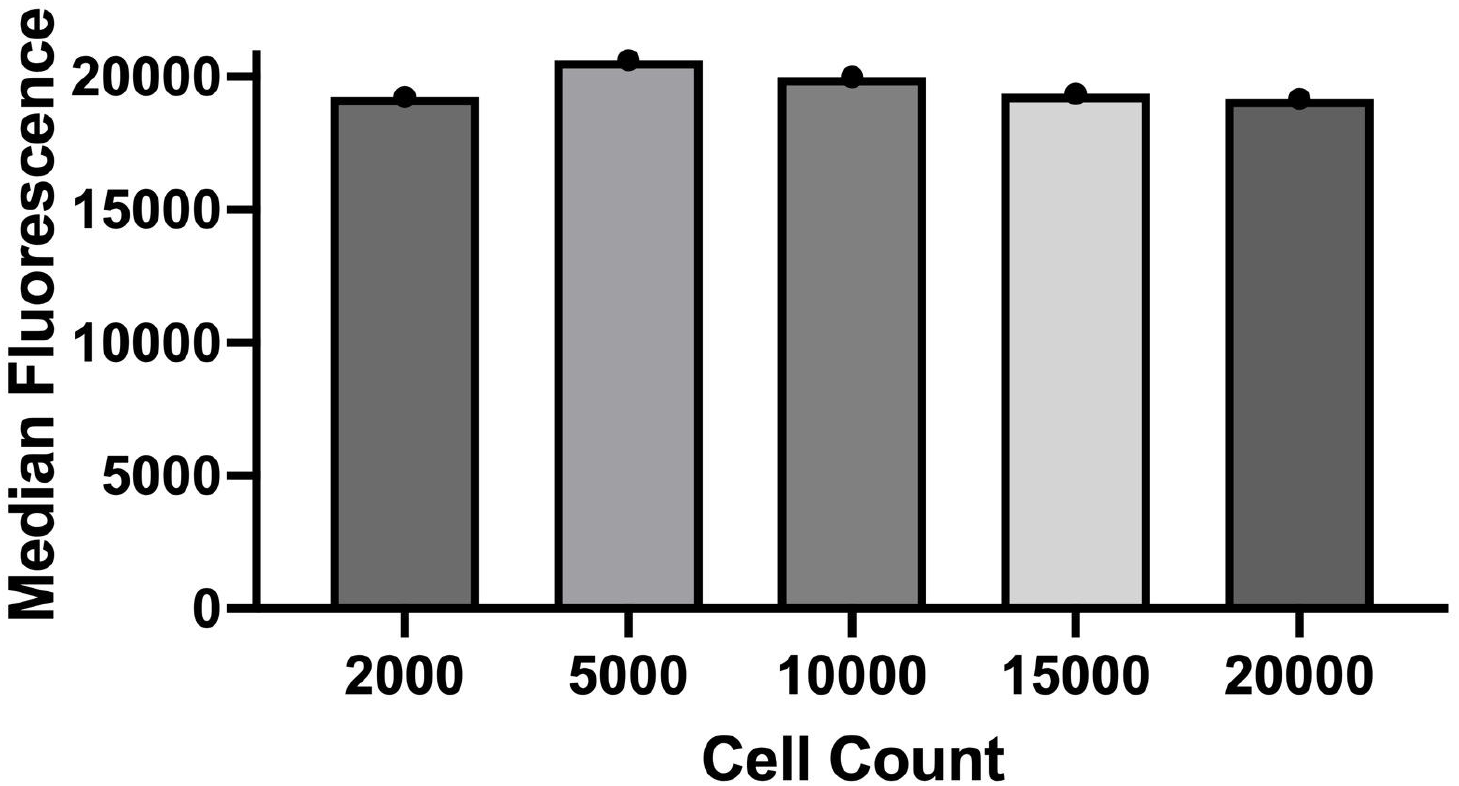
Impact of seeded cell count on fluorescent intensity. A range of U251 cells (from 2000 to 20000) were plated on echistatin conjugated RAD-TGT wells and incubated for 90 minutes. Cells were then dissociated and analyzed the fluorescence intensity with flow cytometery to determine how cell density impacts readout. Fluorescent intensity remained relatively constant regardless of cell seeding density.

**Supplementary Fig. 4.**
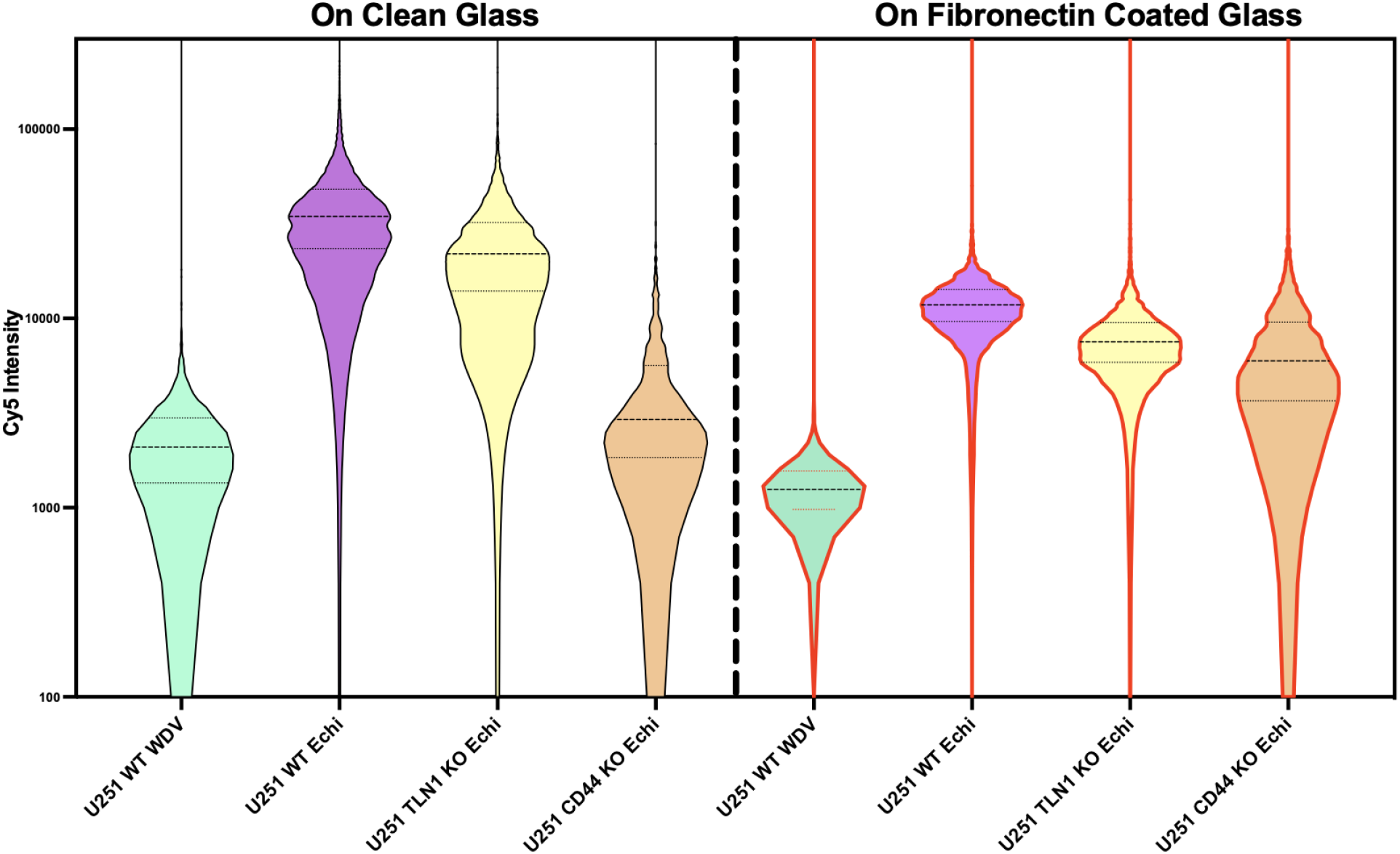
RAD-TGTs Function With and Without Fibronectin Present. RAD-TGTs were plated on both clean glass surfaces and surfaces that were coated with fibronectin. U251 cells were plated on both surfaces and with and without echistatin conjugated to the RAD-TGT, additionally TLN1 KO and CD44 KO U251 cells were tested. Interestingly the same trend was observed between both surfaces with TLN1 KO and CD44 KO having decreased signal relative to WT. Although there were some differences, such as CD44 KO resulting in a less dramatic loss of signal when on fibronectin, we conclude that RAD-TGTs can function in both homogeneous RAD-TGT only surfaces and complex environments such as in conjunction with fibronectin to mimic the extracellular matrix. Differences between the two conditions were attributed to cell behavior changing in a ligand dependent manner and competition between the RAD-TGTs and fibronectin for binding to the cell

**Supplementary Fig. 5.**
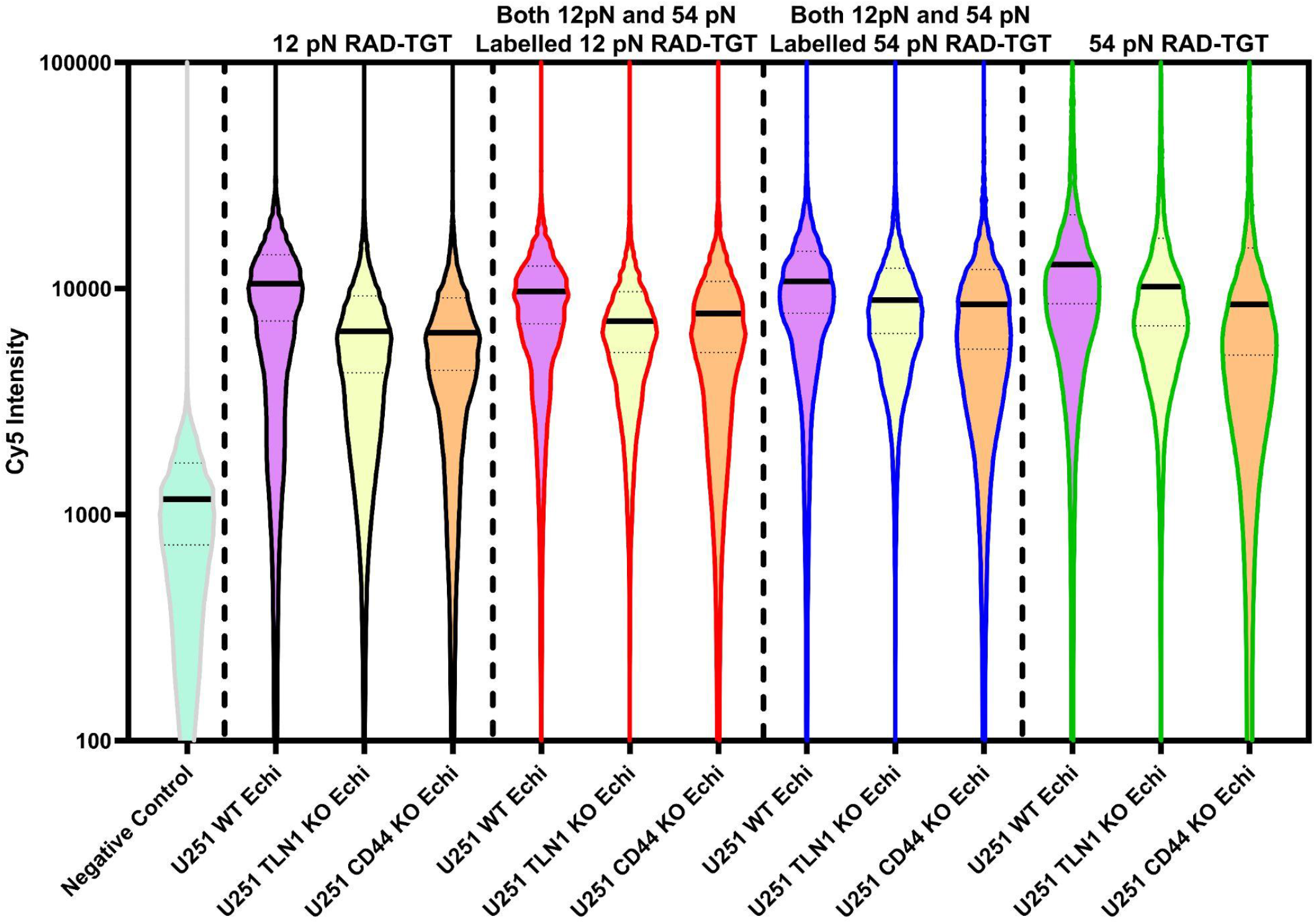
RAD-TGTs retain function with TGTs of different rupture forces and on mixed TGT surfaces. 4 unique RAD-TGTs conjugated to echistatin were compared head to head and to a no-ligand negative control. In all 4 conditions ½ of the RAD-TGTs were labeled with Cy5 and ½ were unlabelled. Condition 1 (black outlines) only contained 12 pN RAD-TGTs. Condition 2 (red outlines) contained both 12 and 54 pN RAD-TGTs but only the 12 pN TGTs were fluorescently labeled, condition 3 (blue outlines) was similar to condition 2 but the 54 pN TGTs were labeled. Condition 4 (green outlines) contained 54 pN RAD-TGTs only. Three variants of U251 cells were plated on these conditions: WT, TLN1 KO, and CD44 KO. Under all conditions the same trend was seen with WT having a higher signal than both knockouts recapitulating the data seen in the main text as indicated by the median fluorescent values (black line in each sample). These results indicate that RAD-TGTs allow for different forces to be tested individually and in conjunction with each other.

**Supplementary Fig. 6.**
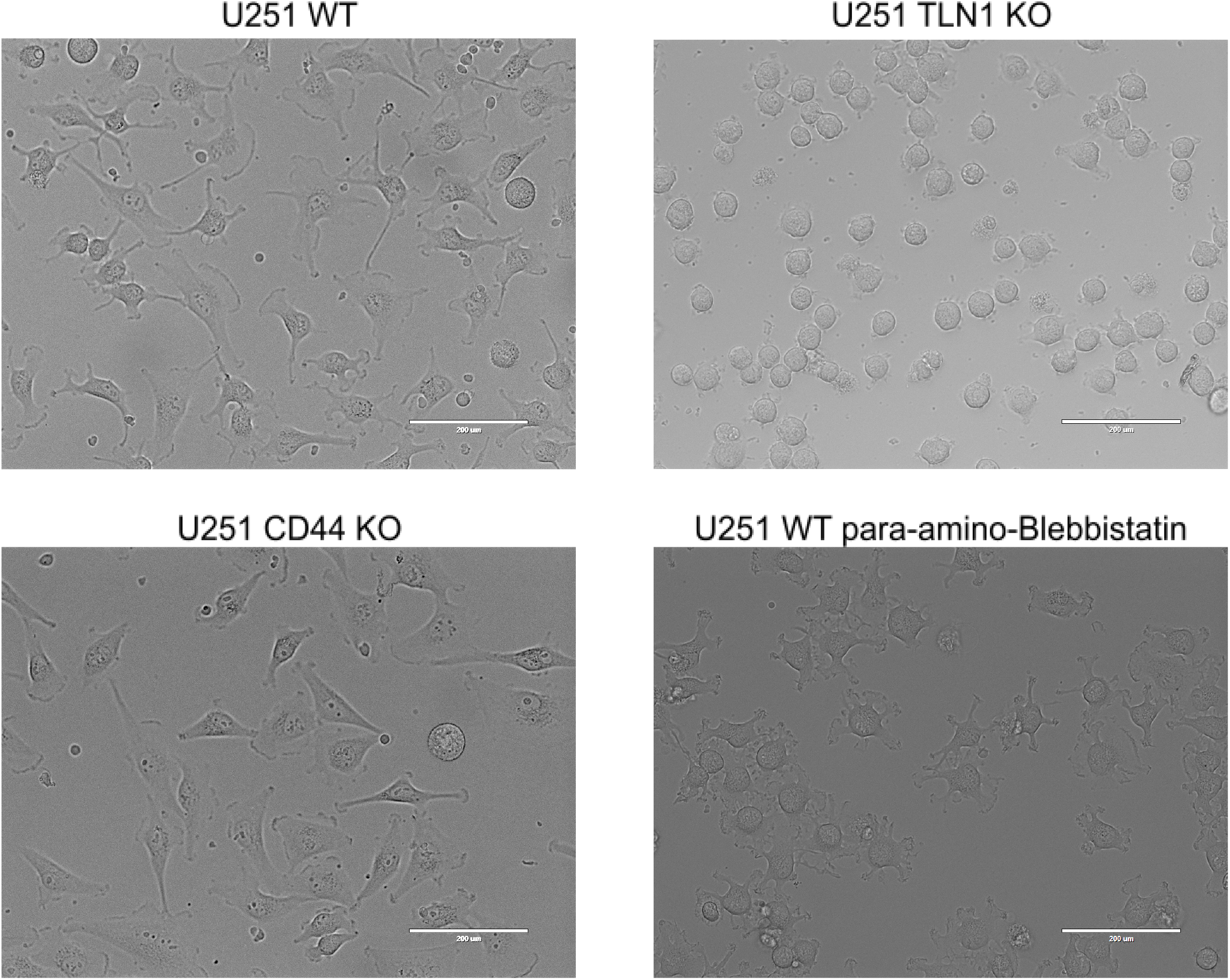
Representative Images of U251 Cells on RAD-TGT surface. All U251 cells and variants were plated on surfaces containing echistatin conjugated RAD-TGTs and allowed to incubate for 90 minutes. In total 4 conditions are shown here, U251 WT, U251 TLN1 KO, U251 CD44 KO, and U251 treated with 50 μM para-amino-Blebbistatin.

**Supplementary Fig. 7.**
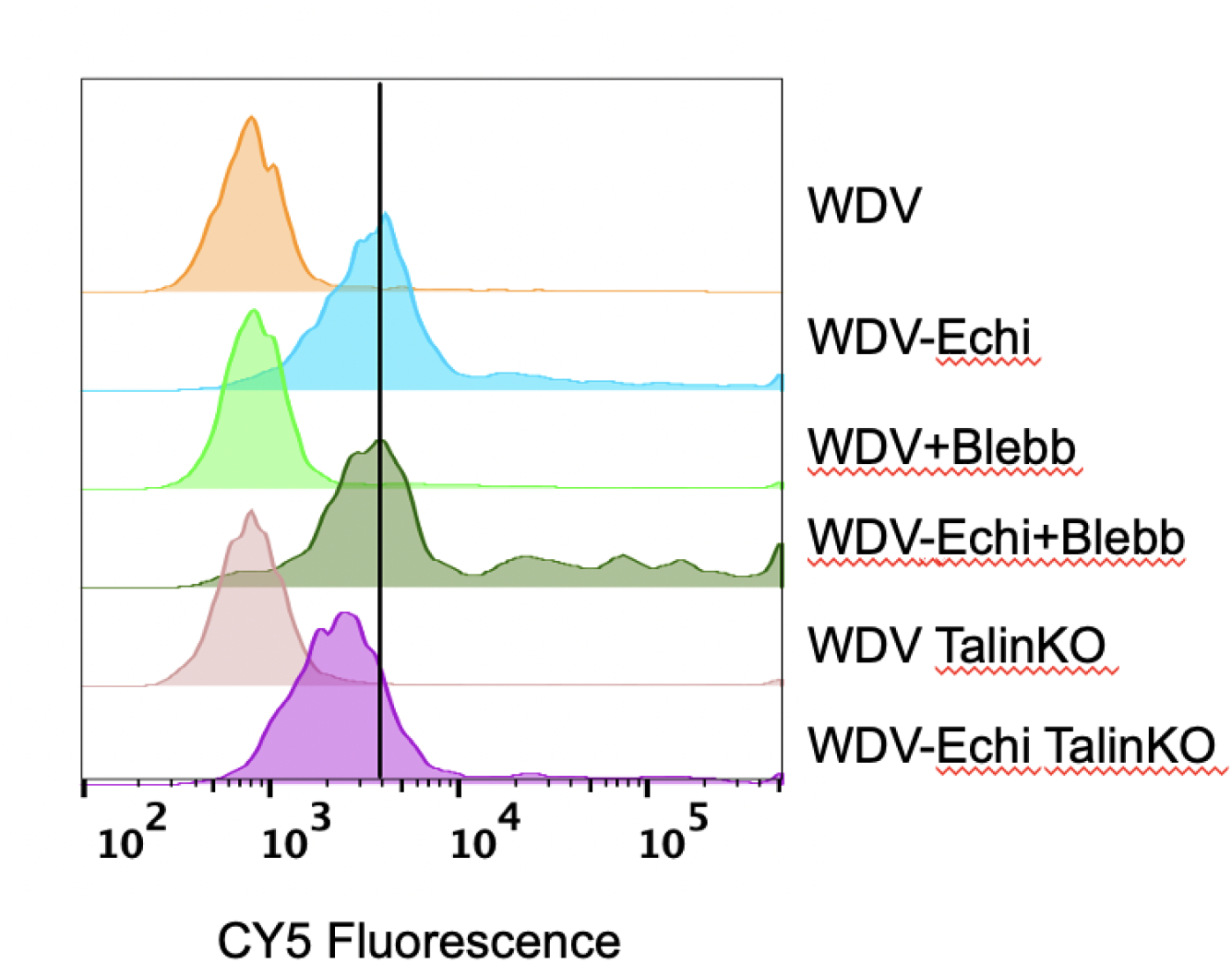
Representative histograms of U251 cells on polystyrene plates. U251 cells were incubated on an echistatin conjugated RAD-TGT surface as described in the main text but rather than glass plates neutravidin coated polystyrene plates (Thermo Fisher, 15129) were used so no biotinylated-BSA or neutravidin was added to the plate. Surfaces contained either no igand WDV conjugated RAD-TGTs or echistatin conjugated RAD-TGTs present. Wild type U251 cells were tested with and without para-amino-blebbistatin treatment and U251 TLN1 KO cells were also measured. The vertical line is representative of the median Cy5 intensity for WDV-Echi. The distribution of events is broad relative to experiments on glass plates and a distinct tail forms in the presence of echistatin. We attributed these results to non-specific binding promoted by the readily adhering polystyrene causing uneven distribution of RAD-TGTs and fluorescently labeled oligos

**Supplementary Fig. 8.**
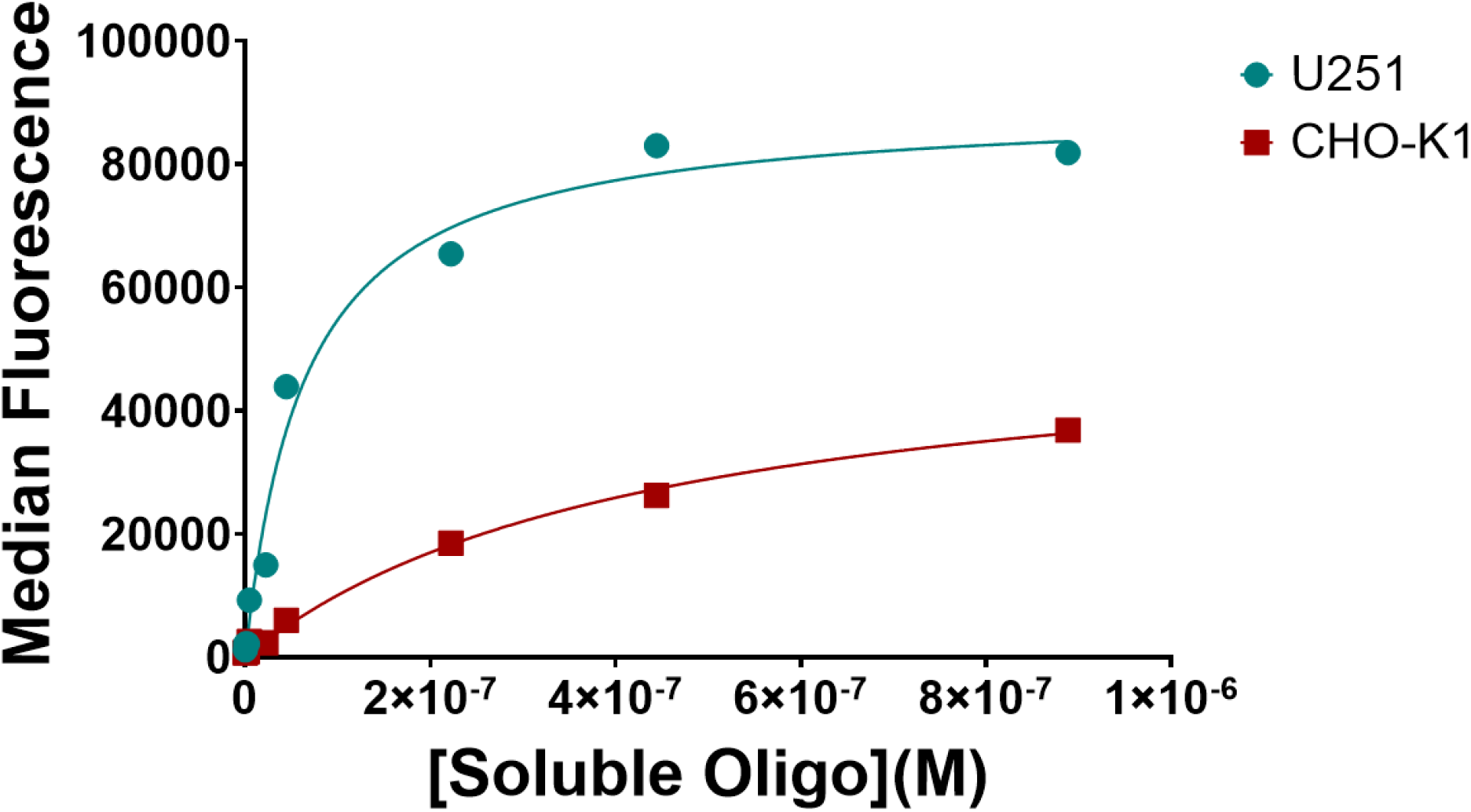
Soluble RAD-TGT Titration Highlights Differences in Integrin Composition of Between Cell Lines. 15,000 U251 and CHO-K1 cells were plated on fibronectin surfaces and allowed to adhere. Soluble RAD-TGTs composed of the echistatin conjugated to fluorescently labeled ligand strand annealed to the non-anchor strand was added into the wells at varying concentrations ranging from 0 to 0.89 μM. The total pmol range of oligo was from 0 to 160 pmol, the standard RAD-TGT experiment in the main text contained 80 pmol oligo assuming all oligo properly adhered, thus this is representative of experimental conditions. From this it is evident that U251 cells internalize the soluble RAD-TGT greater than the CHO-K1 cells,

**Supplementary Fig. 9.**
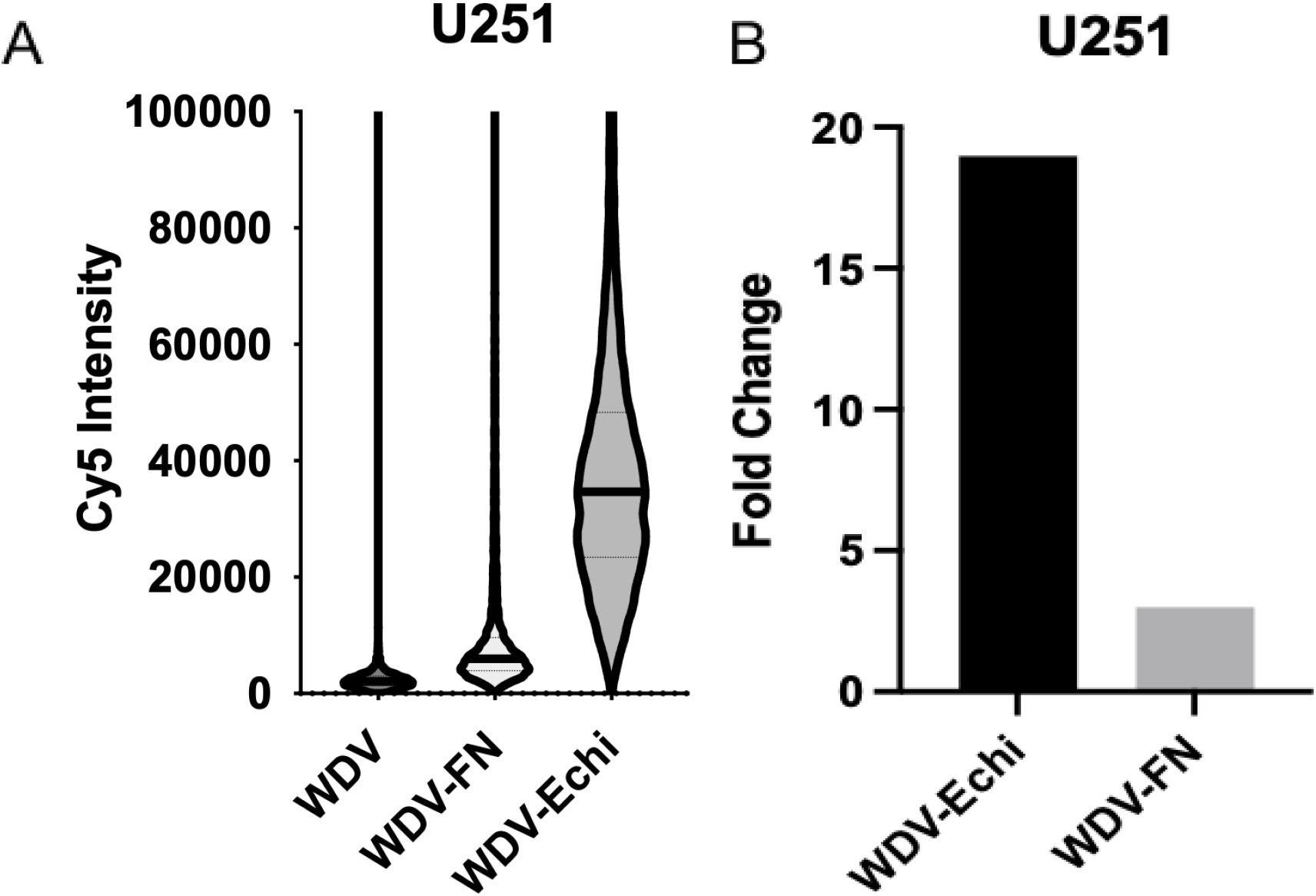
Fluorescent intensity and fold change of RAD-TGTs with different ligands. U251 cells were plated on RAD-TGT surfaces that were either conjugated to no ligand (WDV), a fibronectin derived ligand (FN), or echistatin (Echi). Fluorescence intensity was measured with flow cytometry (A). Median fluorescent value was measured for each condition and fold change was calculated for each ligand by dividing the ligand’s median value by WDV’s value (B). Echi conjugated RAD-TGTs had greater fluorescent signal than FN, this was attributed to echistatin’s high affinity for integrins.

**Supplementary Fig. 10.**
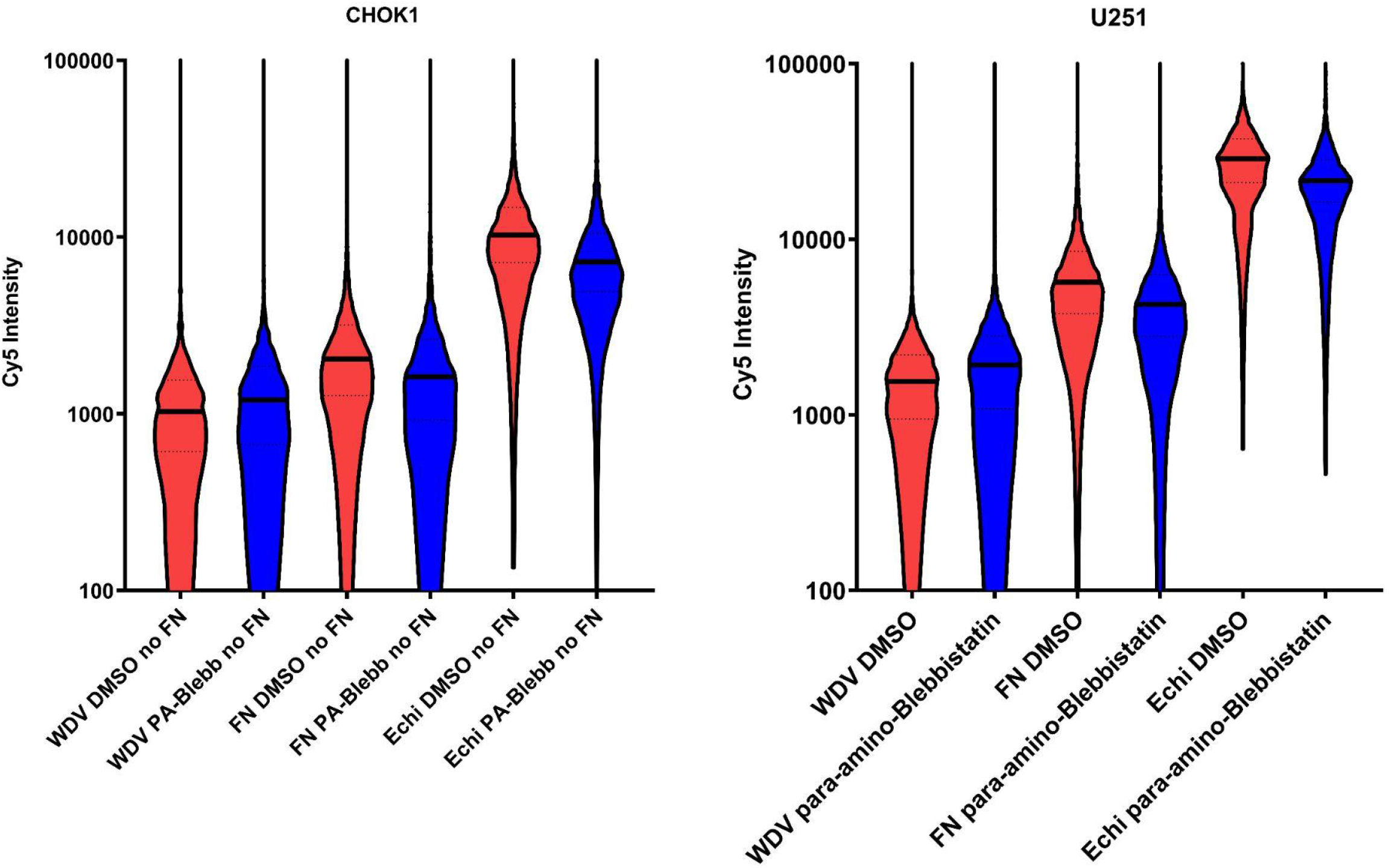
Comparing RAD-TGTs conjugated to different ligands. CHO-K1 cells (left) and U251 cells (right) were plated on Cy5- labeled RAD-TGTs conjugated to WDV, WDV-Fibronectin (FN), or WDV-Echistatin (echi) and treated with DMSO or para-amino-Blebbistatin. Though the fold-change from WDV is smaller for the FN ligand than the Echi ligand, blebbistatin still reduces median fluorescence.

**Supplementary Fig. 11.**
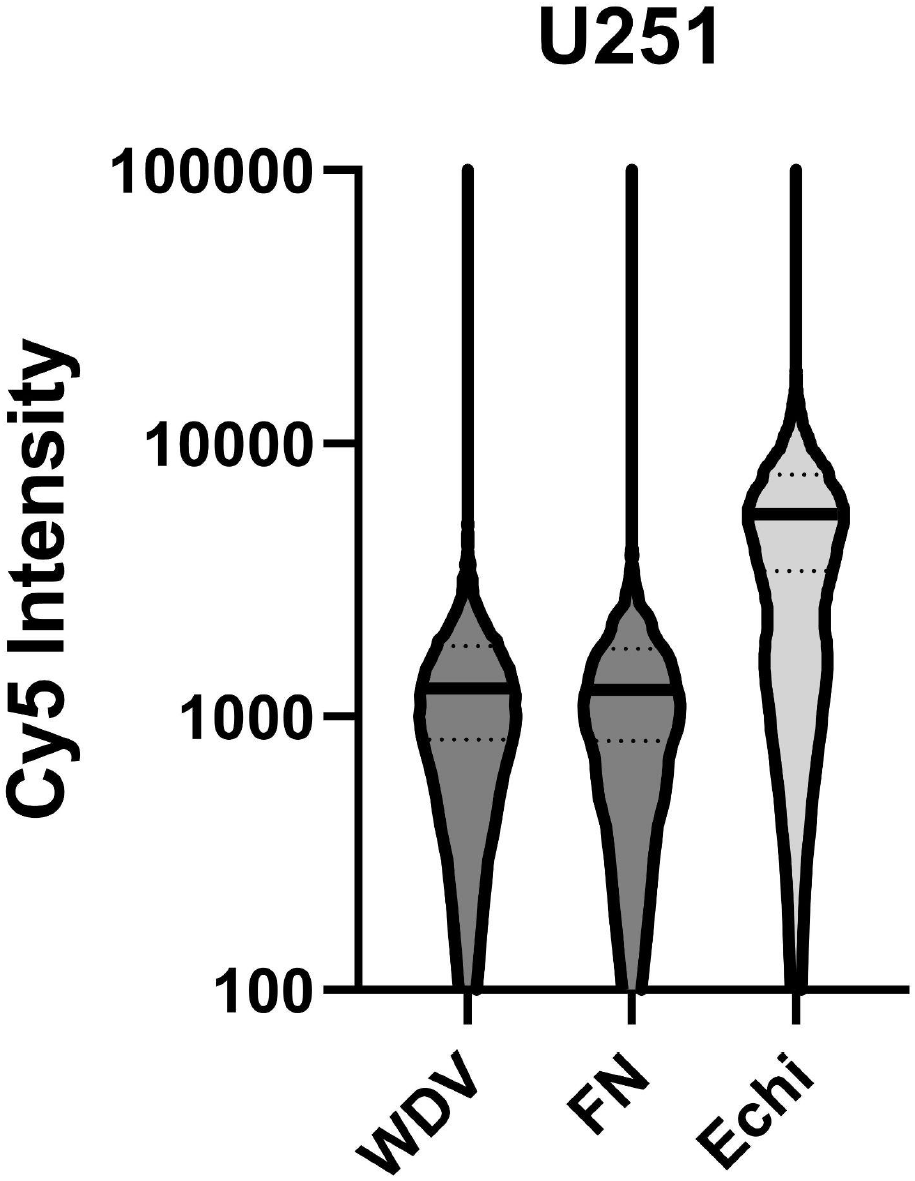
Flow cytometry readout of multiple ligands in one well as done in sequencing experiments. U251 cells were plated in wells containing equimolar amounts of three unique RAD-TGTs one for each ligand (WDV, FN, or Echi) similar to the sequencing experimental setup. Unlike the sequencing experiment one RAD-TGT per well was fluorescently labeled, and three wells were prepared so that each well contained fluorescently labeled RAD-TGTs conjugated to one of the 3 ligands. Each well was then analyzed via flow cytometry and the recorded fluorescence was attributed to the labeled ligand (ie: violin plot labeled FN originated from the well containing fluorescently labeled RAD-TGT conjugated to FN)

**Supplementary Fig. 12.**
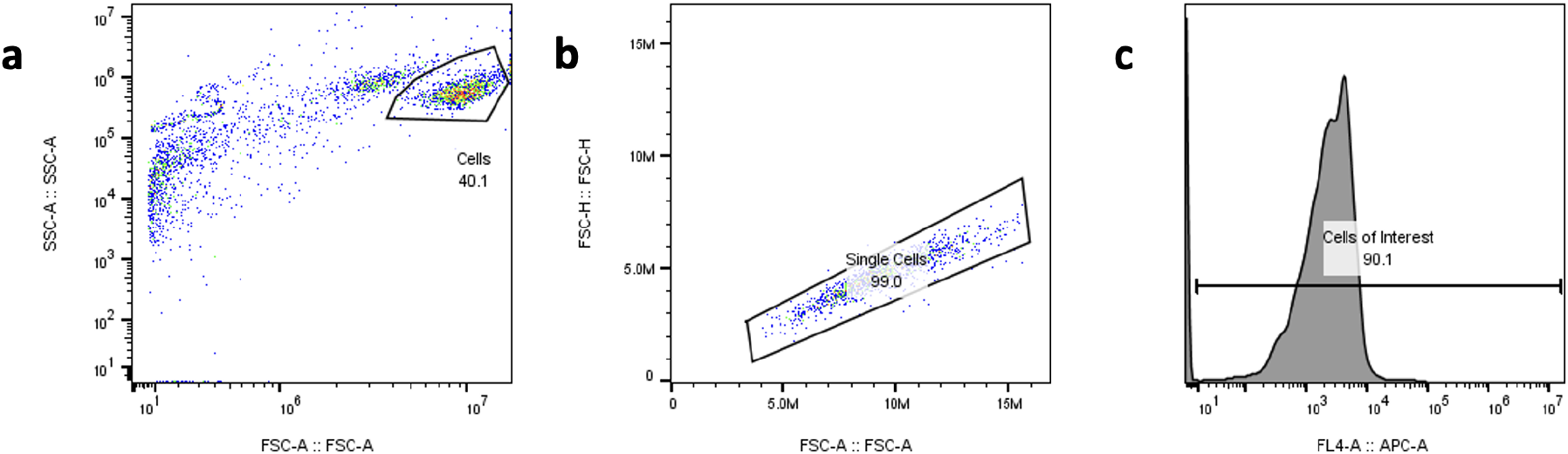
Gating strategy. a. Population of cells was first identified by creating a forward vs side scatter Logicle plot. b. Population identified in a was then then gated to isolate single cells only, this was done by graphing the forward scatter area by height and gating for cells that display a linear relationship. c. Cells from b are then gated to remove any non fluorescent data points. Cells were displayed as a histogram of the fluorescent intensity of the fluorophore of interest and any point with an intensity greater than 10 were collected.

## Notes

### Competing Interest Statement

The authors have declared no competing interest.

## References Cited

1. Reffay, M. et al. Interplay of RhoA and mechanical forces in collective cell migration driven by leader cells. Nat. Cell Biol. 16, 217–223 (2014).

2. Vining, K. H. & Mooney, D. J. Mechanical forces direct stem cell behaviour in development and regeneration. Nat. Rev. Mol. Cell Biol. 18, 728–742 (2017).

3. Ma, V. P.-Y. et al. The magnitude of LFA-1/ICAM-1 forces fine-tune TCR-triggered T cell activation. Science Advances 8, eabg4485 (2022).

4. Acerbi, I. et al. Human breast cancer invasion and aggression correlates with ECM stiffening and immune cell infiltration. Integr. Biol. 7, 1120–1134 (2015).

5. Fenner, J. et al. Macroscopic stiffness of breast tumors predicts metastasis. Sci. Rep. 4, 5512 (2014).

6. Zuela-Sopilniak, N. & Lammerding, J. Can’t handle the stress? Mechanobiology and disease. Trends Mol. Med. (2022) doi:10.1016/j.molmed.2022.05.010.

7. Sheridan, C. Pancreatic cancer provides testbed for first mechanotherapeutics. Nat. Biotechnol. 37, 829–831 (2019).

8. Fischer, L. S., Rangarajan, S., Sadhanasatish, T. & Grashoff, C. Molecular Force Measurement with Tension Sensors. Annu. Rev. Biophys. 50, 595–616 (2021).

9. Ham, T. R., Collins, K. L. & Hoffman, B. D. Molecular Tension Sensors: Moving Beyond Force. Curr Opin Biomed Eng 12, 83–94 (2019).

10. Grashoff, C. et al. Measuring mechanical tension across vinculin reveals regulation of focal adhesion dynamics. Nature 466, 263–266 (2010).

11. LaCroix, A. S., Lynch, A. D., Berginski, M. E. & Hoffman, B. D. Tunable molecular tension sensors reveal extension-based control of vinculin loading. Elife 7, (2018).

12. Aird, E. J., Tompkins, K. J., Ramirez, M. P. & Gordon, W. R. Enhanced Molecular Tension Sensor Based on Bioluminescence Resonance Energy Transfer (BRET). ACS Sensors (2020) doi:10.1021/acssensors.9b00796.

13. Borghi, N. et al. E-cadherin is under constitutive actomyosin-generated tension that is increased at cell-cell contacts upon externally applied stretch. Proc. Natl. Acad. Sci. U. S. A. 109, 12568–12573 (2012).

14. Zhang, Y., Ge, C., Zhu, C. & Salaita, K. DNA-based digital tension probes reveal integrin forces during early cell adhesion. Nat. Commun. 5, 5167 (2014).

15. Blakely, B. L. et al. A DNA-based molecular probe for optically reporting cellular traction forces. Nat. Methods 11, 1229–1232 (2014).

16. Wang, X. & Ha, T. Defining single molecular forces required to activate integrin and notch signaling. Science 340, 991–994 (2013).

17. Sarkar, A., Zhao, Y., Wang, Y. & Wang, X. Force-activatable coating enables high-resolution cellular force imaging directly on regular cell culture surfaces. Phys. Biol. 15, 065002 (2018).

18. Morimatsu, M., Mekhdjian, A. H., Adhikari, A. S. & Dunn, A. R. Molecular tension sensors report forces generated by single integrin molecules in living cells. Nano Lett. 13, 3985–3989 (2013).

19. Zhao, B. et al. Quantifying tensile forces at cell-cell junctions with a DNA-based fluorescent probe. Chem. Sci. 11, 8558–8566 (2020).

20. Paszek, M. J. et al. The cancer glycocalyx mechanically primes integrin-mediated growth and survival. Nature 511, 319–325 (2014).

21. Deal, B. R., Brockman, J. M. & Salaita, K. DNA probes that store mechanical information reveal transient piconewton forces applied by T cells. Proceedings of the (2019).

22. Haas, A. J. et al. Interplay between Extracellular Matrix Stiffness and JAM-A Regulates Mechanical Load on ZO-1 and Tight Junction Assembly. Cell Rep. 32, 107924 (2020).

23. Ramirez, M. P. et al. Dystrophin missense mutations alter focal adhesion tension and mechanotransduction. Proc. Natl. Acad. Sci. U. S. A. 119, e2205536119 (2022).

24. Hu, Y. et al. DNA-based microparticle tension sensors (μTS) for measuring cell mechanics in non-planar geometries and for high-throughput quantification. Angew. Chem. Weinheim Bergstr. Ger. 133, 18192–18198 (2021).

25. Lovendahl, K. N., Hayward, A. N. & Gordon, W. R. Sequence-Directed Covalent Protein-DNA Linkages in a Single Step Using HUH-Tags. J. Am. Chem. Soc. 139, 7030–7035 (2017).

26. Wang, X. et al. Constructing modular and universal single molecule tension sensor using protein G to study mechano-sensitive receptors. Sci. Rep. 6, 21584 (2016).

27. Kapp, T. G. et al. A Comprehensive Evaluation of the Activity and Selectivity Profile of Ligands for RGD-binding Integrins. Sci. Rep. 7, 39805 (2017).

28. Duan, Y. et al. Mechanically Triggered Hybridization Chain Reaction. Angewandte Chemie vol. 133 20127–20134 (2021).

29. Tompkins, K. J. et al. Molecular underpinnings of ssDNA specificity by Rep HUH-endonucleases and implications for HUH-tag multiplexing and engineering. Nucleic Acids Res. (2021) doi:10.1093/nar/gkaa1248.

30. Everett, B. A. et al. Crystal structure of the Wheat dwarf virus Rep domain. Acta Crystallogr. Sect. F Struct. Biol. Cryst. Commun. 75, 744–749 (2019).

31. Roein-Peikar, M., Xu, Q., Wang, X. & Ha, T. Ultrasensitivity of Cell Adhesion to the Presence of Mechanically Strong Ligands. Phys. Rev. X 6, 011001 (2016).

32. Jo, M. H. et al. Molecular Nanomechanical Mapping of Histamine-Induced Smooth Muscle Cell Contraction and Shortening. ACS Nano (2021) doi:10.1021/acsnano.1c01782.

33. Bangasser, B. L. et al. Shifting the optimal stiffness for cell migration. Nat. Commun. 8, 15313 (2017).

34. Kim, Y. & Kumar, S. CD44-mediated adhesion to hyaluronic acid contributes to mechanosensing and invasive motility. Mol. Cancer Res. 12, 1416–1429 (2014).

35. Roy, D. C., Wilke-Mounts, S. J. & Hocking, D. C. Chimeric fibronectin matrix mimetic as a functional growth-and migration-promoting adhesive substrate. Biomaterials 32, 2077–2087 (2011).

36. Sanders, H. M. H. F. et al. The binding of CNA35 contrast agents to collagen fibrils. Chem. Commun. 47, 1503–1505 (2011).

37. Marko, T. A. et al. Slit-Robo GTPase-Activating Protein 2 as a metastasis suppressor in osteosarcoma. Sci. Rep. 6, 39059 (2016).

38. Moriarity, B. S. et al. Simple and efficient methods for enrichment and isolation of endonuclease modified cells. PLoS One 9, e96114 (2014).

39. Kluesner, M. G. et al. EditR: A Method to Quantify Base Editing from Sanger Sequencing. CRISPR J 1, 239–250 (2018).

40. Rausch, T., Fritz, M. H.-Y., Untergasser, A. & Benes, V. Tracy: basecalling, alignment, assembly and deconvolution of sanger chromatogram trace files. BMC Genomics 21, 230 (2020).

41. Cock, P. J. A. et al. Biopython: freely available Python tools for computational molecular biology and bioinformatics. Bioinformatics 25, 1422–1423 (2009).

42. Lord, S. J., Velle, K. B., Mullins, R. D. & Fritz-Laylin, L. K. SuperPlots: Communicating reproducibility and variability in cell biology. J. Cell Biol. 219, (2020).

